# Programmable RNA detection generates DNA barcodes for multiplexed phage-host interaction screening

**DOI:** 10.64898/2026.02.13.705812

**Authors:** Jihoon Han, Seth L. Shipman

## Abstract

Here, we introduce Detectrons, modular biosensors that couples programmable toehold switches with retron-mediated reverse transcription to transduce RNA inputs into unique DNA barcodes. The ability to convert dynamic RNA signals into durable DNA records within living cells unlocks powerful new modes of transcript-based sensing with applications including viral infection detection. Through the construction of a synthetic toehold–retron library and application of machine learning, we uncovered key design principles that improve signal strength and specificity. We applied Detectrons to the multiplexed live-cell detection of specific phage infections, enabling transcript-triggered barcode synthesis and quantitative host susceptibility profiling in pooled bacterial populations. Detectrons are the first RNA-to-DNA transduction system, directly linking transient RNA detection to stable, sequence-encoded DNA outputs. This platform provides a scalable and generalizable strategy for phage screening and for recording transcriptional events in complex bacterial communities.

## INTRODUCTION

The rapid escalation of antimicrobial resistance (AMR) has reestablished bacterial infections as a significant global health threat. By 2050, AMR-induced mortality is projected to reach 10 million deaths per year, surpassing mortality from diseases such as cancer and diabetes^*1*^. This alarming trend underscores the urgent need for innovative therapeutic strategies for bacterial infections. Bacteriophages, the natural predators of bacteria, offer a promising solution to managing pathogenic bacteria, especially as potential alternatives to antibiotics^*2*^. The high selectivity for specific bacterial species allows phage to eliminate the intended pathogens without disrupting the broader microbiota–a significant advantage over broad-spectrum antibiotics that can harm beneficial bacteria^*3*^. However, this specificity also introduces a major challenge in the efficient identification and screening of phages with therapeutic potential against specific bacterial strains^*4*^. Traditional methods, like plaque assays, are time-consuming, labor-intensive, and limited in throughput, often requiring extensive resources, which delays timely interventions. Thus, what we need is a high-throughput, multiplexable screening platform that can efficiently determine phage host range in short time within a single-pot setup.

Synthetic biology offers transformative solutions to such challenges by repurposing and engineering biological components for practical applications^5–8^. One notable innovation is the programmable RNA sensors, known as toehold switches. These sensors can be rationally designed to specifically bind and detect virtually any RNA sequence, offering unparalleled orthogonality, programmability, and precision^*9,10*^. Such advances have successfully facilitated the development of diagnostic tools aimed at combating global health crises^*11,12*^. However, the utility of toehold switches in multiplexing and high-throughput applications, such as screening numerous RNAs in a single-pot reaction, is constrained by their reliance on protein-based output signals. While protein outputs can be useful for real-time and functional readouts, they significantly lack the diversity, sensitivity, programmability, and multiplexing capacity of DNA outputs, which are more stable and better suited for biosensors and synthetic circuits.

To address this limitation, we developed Detectrons, a multiplexable biosensor that detects RNA and generates DNA as output signal (**Fig. 1a**). We employ retrons—bacterial genetic elements that produce multicopy single-stranded DNA (msDNA) using a reverse transcriptase (RT) and a noncoding RNA (ncRNA) template—as the output generator ^*13,14*^. Retrons, which are immune systems in bacteria, have been harnessed as intracellular factories for single-stranded DNA (ssDNA) in various biotechnological applications, including genome editing where the natural ncRNA template is partially replaced with a desired sequence such as an editing donor^*15– 20*^. In our approach, distinct DNA barcodes are linked to specific toehold switches (**Fig. 1a**). In the absence of the target RNA that the toehold is designed to detect, the toehold switch inhibits retron-mediated DNA barcode production by sequestering a critical component of the retron that is necessary for reverse transcription. Upon detection of the target RNA, the toehold switch undergoes strand-displacement, triggering retron-mediated DNA barcode synthesis (**Fig. 1b**). This flexible new platform has many potential applications. In this work, we linked distinct phage-sensing modules to orthogonal barcode outputs. Detectrons enable digital, multiplexed detection of phage infections in a pooled culture, and through dual barcoding, resolve strain-specific susceptibilities within mixed populations—allowing scalable, high-throughput profiling of phage– host interactions.

**Figure 1.**
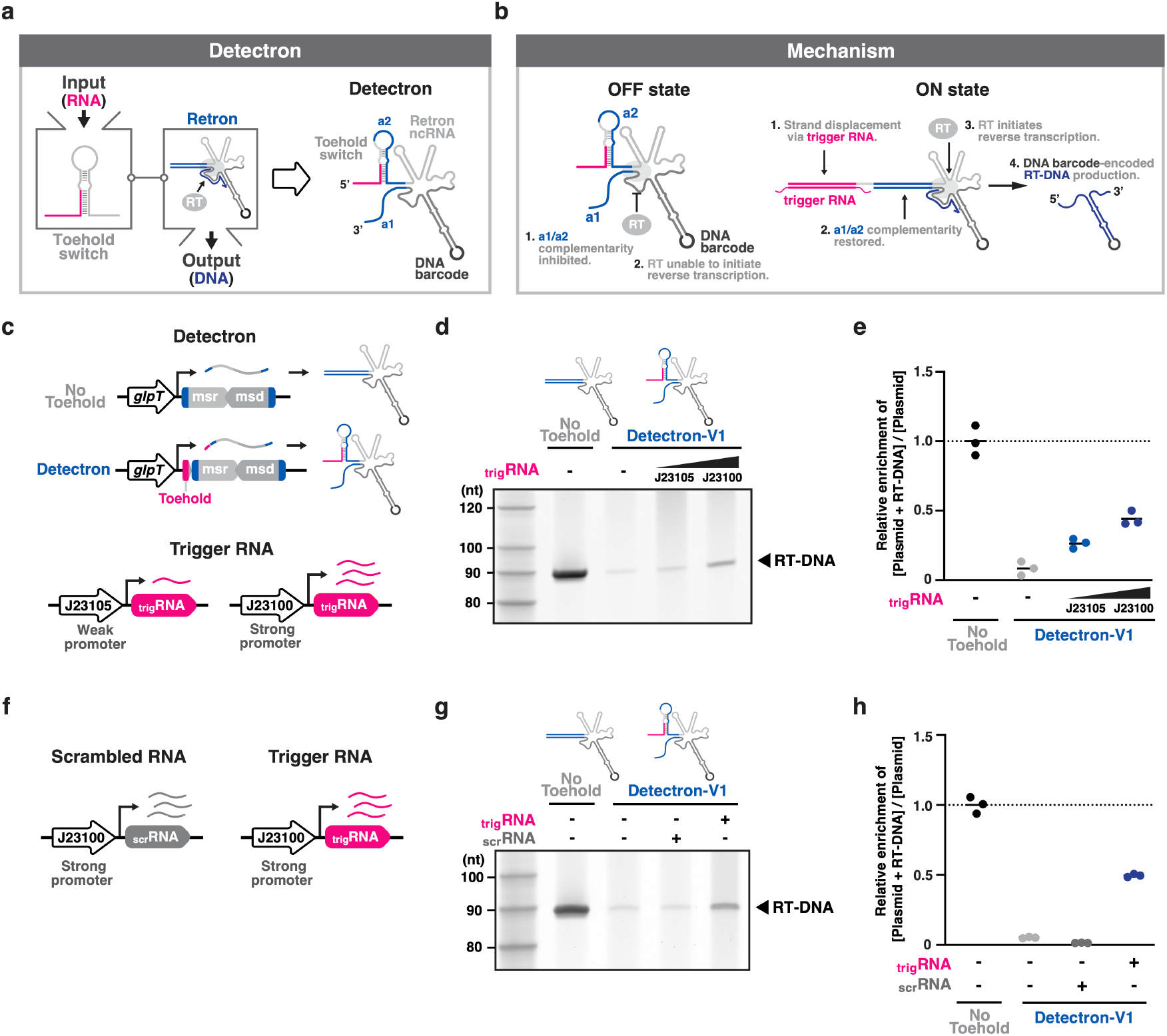
Detectrons specifically report on target RNA by DNA barcode synthesis. **a**, Schematic of the biological components of a Detectron. RT, reverse transcriptase. **b**, Schematic of the Detectron mechanism. **c**, Schematic of the experimental promoters used to express No Toehold control, Detectron, and trigger RNA, along with representations of the expressed transcripts. msr, multicopy single-stranded RNA. msd, multicopy single-stranded DNA. trigRNA, trigger RNA. **d**, Urea PAGE analysis of RT-DNA from the No Toehold control (lane 1, excluding ladder) and Detectron in the absence (lane 2) or presence of trigger RNA expressed under the J23105 promoter (lane 3) or J23100 promoter (lane 4). **e**, Enrichment of RT-DNA + plasmid over plasmid alone relative to No Toehold control, as measured by qPCR; One-way ANOVA with Tukey’s multiple comparisons test (corrected): Detectron versus No Toehold: *P* < 0.0001; Detectron + trigger RNA (J23105) versus Detectron: *P* = 0.0439; Detectron + trigger RNA (J23100) versus Detectron: *P* = 0.0008; Detectron + trigger RNA (J23100) versus Detectron + trigger RNA (J23105): *P* = 0.0481. Closed circles indicate three biological replicates. Bars represent the mean of three biological replicates. **f**, Schematic of the experimental promoters used to express scrambled RNA and trigger RNA. scrRNA, scrambled RNA. **g**, Urea PAGE analysis of RT-DNA from the No Toehold control (lane 1) and Detectron in the absence (lane 2) or presence of scrambled RNA (lane 3) or trigger RNA (lane 4). **h**, Enrichment of RT-DNA + plasmid over plasmid alone relative to No Toehold, as measured by qPCR; One-way ANOVA with Tukey’s multiple comparisons test (corrected): Detectron versus No Toehold: *P* < 0.0001; Detectron + trigger RNA (J23100) versus Detectron: *P* < 0.0001; Detectron + trigger RNA (J23100) versus Detectron + scrambled RNA (J23100): *P* < 0.0001. Closed circles indicate three biological replicates. Bars represent the mean of three biological replicates.

## RESULTS

### Detectrons specifically report on target RNA by DNA barcode synthesis

The retron ncRNA consists of two distinct, partially overlapping regions: the multicopy single-stranded RNA (msr), which is not reverse transcribed, and the multicopy single-stranded DNA (msd), which undergoes reverse transcription. The 5’ (a2) and 3’ (a1) ends of the ncRNA are complementary and fold back on themselves, forming a structural element known as a1/a2, which is essential for initiating reverse transcription at a conserved priming guanosine (**Extended Data Fig. 1**). We previously demonstrated that reducing a1/a2 complementarity below 7 base pairs (bp) substantially impairs reverse transcribed DNA (RT-DNA) production^*18*^. Based on this, we hypothesized that sequestering the majority of the a2 region within a toehold switch hairpin would disrupt a1/a2 complementarity and inhibit RT-DNA synthesis. In the presence of target RNA, the toehold switch will undergo strand-displacement, restoring a1/a2 complementarity and initiating reverse transcription (**Fig. 1b**).

To test our design, we utilized a previously developed Zika virus RNA-detecting toehold switch^*11*^ and retron-Eco1 (formerly known as ec86), the first characterized retrons in *Escherichia coli* (*E. coli*)^*21*^ (**Extended Data Fig. 1**). To preserve the specificity and sensitivity of the predesigned toehold switch, we maintained most of its structure and sequence, replacing only the ribosome binding site (RBS) in the loop with a non-RBS sequence to prevent unintended translation of the retron ncRNA. We then repurposed the sequence from the non-RBS to the linker as the a2 region of Eco1 ncRNA, redesigning the a1 sequence accordingly (**Extended Data Fig. 1**). We termed this toehold switch-integrated retron ncRNA a Detectron. When expressed under *glpT* promoter in *E. coli* (bMS.346), Detectron substantially inhibited RT-DNA production, which was partially restored in a dose-dependent manner upon trigger RNA expression, as visualized via polyacrylamide gel electrophoresis (PAGE) and quantified by quantitative polymerase chain reaction (qPCR) (**Fig. 1c-e)**. This restoration did not occur with scrambled RNA expression, indicating that the Detectron is specific for the designed trigger RNA (**Fig. 1f-h**). Similar results were observed when trigger RNA and scrambled RNA were expressed under an inducible promoter (**Extended Data Fig. 2a,b**).

### Optimization of Detectrons using a high-throughput variant library

Although the initial Detectron design was functional, the dynamic range was relatively limited. To optimize the conformational dynamics of Detectrons and enhance their responsiveness to target RNA, we constructed a library of 7,840 Detectron variants by systematically varying each component of the toehold switch—length of stem1, bulge, stem2, and loop—as well as the distance from the stem base to the priming guanosine (d) in the retron ncRNA (**Fig. 2a**). For high-throughput analysis, a unique DNA barcode was encoded into the msd loop of each variant (**Fig. 2b**). By sequencing barcode-encoded RT-DNAs in the absence (OFF state) or presence (ON state) of trigger RNA, we measured the ON/OFF ratios for each variant and identified those with enhanced ON/OFF performance (**Extended Data Fig. 3a**).

**Figure 2.**
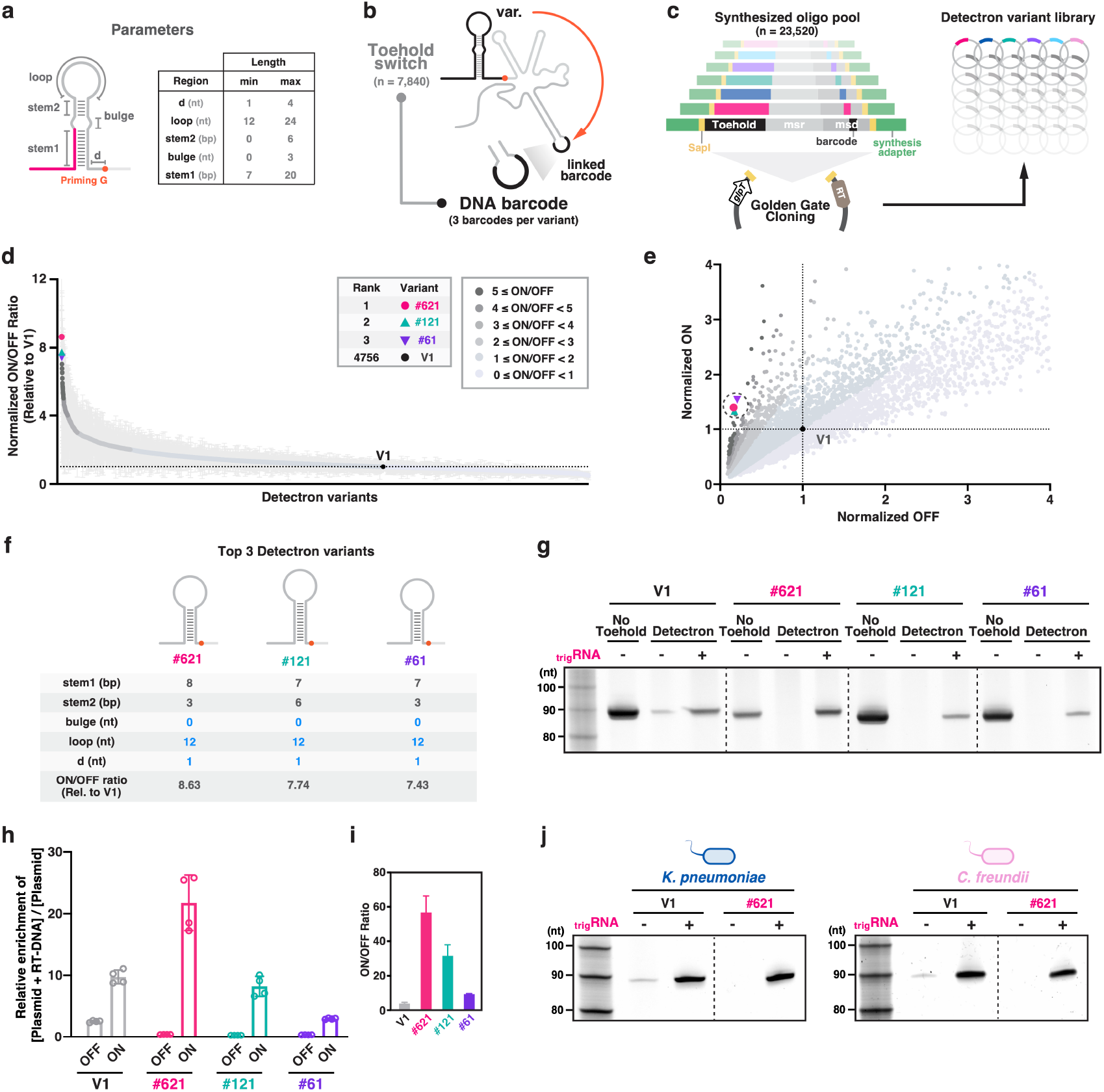
Optimization of Detectrons using a high-throughput variant library. **a**, Schematic of the Detectron variant library design. stem1, stem2, bulge, loop, and the distance from stem base to the priming guanosine (d) were systematically varied to construct 7,840 variants. **b**, Schematic of a barcoded-Detectron variant: each variant was linked to three unique DNA barcodes encoded in the msd loop, expanding the library to 23,520 constructs. **c**, Design of synthesized oligos and Golden Gate cloning workflow for constructing the variant library. **d**, Distribution of ON/OFF ratios across the variant library, ranked from highest to lowest ON/OFF ratio. Each circle represents a Detectron variant, color-coded by the average ON/OFF ratios (±s.d.) of three biological replicates. V1, original Detectron design. **e**, Scatter plot of ON-state versus OFF-state signals across all Detectron variants. Points are color-coded by ON/OFF ratio. **f**, Structural features and parameter values of the top three Detectron variants with the highest the average ON/OFF ratios. **g**, Urea PAGE visualization of RT-DNA from No Toehold control (lane 1, 4, 7, 10, excluding ladder) and the indicated Detectron variants in the absence (lane 2, 5, 8, 11) or presence of trigger RNA expressed under the J23100 promoter (lane 3, 6, 9, 12). **h**, Enrichment of RT-DNA + plasmid over plasmid alone, as measured by qPCR; One-way ANOVA with Tukey’s multiple comparisons test (corrected) for OFF-state signal: #621 OFF versus V1 OFF: *P* < 0.0001; #121 OFF versus V1 OFF: *P* < 0.0001; #61 OFF versus V1 OFF: *P* < 0.0001; One-way ANOVA with Tukey’s multiple comparisons test (corrected) for ON-state signal: #621 ON versus V1 ON: *P* < 0.0001. Closed circles indicate four biological replicates. Bars represent the mean (±s.d.) of four biological replicates. **i**, ON/OFF ratios of indicated Detectron variants; One-way ANOVA with Tukey’s multiple comparisons test (corrected): #621 versus V1: *P* < 0.0001; #121 versus V1: *P* < 0.0001. Bars represent the mean (±s.d.) of four biological replicates. **j**, Urea PAGE visualization of RT-DNA from V1 and #621 Detectrons in the absence (lane 1, 3) or presence of trigger RNA expressed under the J23100 promoter (lane 2, 4) in *K. pneumoniae* (left) and *C. freundii* (right).

To ensure accurate quantification of RT-DNA levels corresponding to each configuration while minimizing the potential impact of specific barcode sequences on RT-DNA production, each Detectron variant was linked to three distinct DNA barcodes, generating three sublibraries and expanding the total library size to 23,520 unique constructs (**Fig. 2b** and **Extended Data Fig. 3b**). For reliable barcode differentiation in high-throughput sequencing, we designed 13-nt DNA barcodes with a minimum Hamming distance of 4, reducing the risk of barcode misclassification due to sequencing errors while enabling robust multiplexing. Additionally, the GC content was constrained between 30% and 70% to enhance sequence stability, minimize secondary structure formation, and ensure efficient amplification and accurate readout (**Extended Data Fig. 3c**). Complete variant and barcode sequences are provided in **Supplementary Tables S4**. To circumvent current practical limits of 300 bp on oligo pool synthesis, we first generated a minimal Eco1 ncRNA by deleting 12 bp from the P4 stem (**Extended Data Fig. 4a**). Although this variant produced less RT-DNA than the wild-type (wt), it remained detectable and functional in our system (**Extended Data Fig. 4b,c**). Despite using the minimal Eco1 ncRNA, a subset of oligos exceeded the 300 bp limit for oligo pool synthesis. To address this, we removed the invariable region—comprising the msr and the first 29 bp of msd—and inserted a Type IIS restriction enzyme site (BsaI) to enable efficient assembly (**Extended Data Fig. 4d)**. We then implemented a two-step Golden Gate cloning strategy: in the first step, synthesized oligos were cloned into the backbone; in the second step, the invariable region was reinserted, restoring the complete Detectron construct (**Extended Data Fig. 4e**).

We constructed two libraries: an ON library (constitutively expressing trigger RNA) and an OFF library (no trigger RNA expression) (**Fig. 2c** and **Extended Data Fig. 3a**). These libraries were transformed into an *E. cloni* 10G strain for RT-DNA production. Three separate transformations of each library serve as biological replicates. After overnight culture, barcode-encoded RT-DNAs along with plasmids were extracted using the QIAGEN Midiprep Plasmid Plus Kit. A portion of the extracted material was reserved for plasmid quantification, and the remainder was subjected to plasmid depletion using the Zymo ssDNA Clean & Concentrator Kit to enrich for single-stranded RT-DNA prior to downstream quantification. Using a previously described sequencing pipeline^16–18,22^, two rounds of PCR amplification were performed to add sequencing adapters and indexes to the extracted RT-DNAs and plasmids, and the resulting products were sequenced on an Illumina platform. The abundance of RT-DNAs from each variant was normalized to the abundance of its corresponding plasmid, which were prepped and sequenced in parallel. As a predefined variance-based exclusion criterion, variants with a standard error of the mean (SEM) greater than 50% of the corresponding mean across replicates (26 of 7,840) were excluded, leaving 7,814 variants for analysis, of which 4,755 exhibited an improved ON/OFF ratio compared to the original Detectron design (V1) (**Fig. 2d**). Most of the variants with high ON/OFF ratios were characterized by a substantial reduction in OFF-state signal and a moderate increase in ON-state signal (**Fig. 2e**). We individually validated the three variants with the highest ON/OFF ratios (#621, #121, and #61), which share common structural characteristics: (i) loop length of 12 bp, (ii) a distance of 1 bp between the stem base and priming G, and (iii) no bulge (**Fig. 2f**). The total stem lengths for these variants were 11, 13, and 10 bp, respectively. We measured individual ON/OFF ratios of these variants using the full-length Eco1 ncRNA. All three variants exhibited a substantially reduced OFF-state signal compared to V1 (**Fig. 2g,h**). Notably, variant #621 showed a significantly enhanced ON-state signal, leading to a 14.7-fold increase in the ON/OFF ratio relative to V1 (**Fig. 2h,i**). Because portability is key for practical deployment, we next asked whether the optimized variant would retain its advantage in other bacterial hosts. Indeed, #621 consistently outperformed V1 in *K. pneumoniae* and *C. freundii* (**Fig. 2j**), demonstrating that the improvements extend beyond *E. coli* and highlighting the robustness of the Detectron platform.

### Machine learning on Detectron variant library results identifies design principles for optimal ON/OFF ratios

Having identified the tested Detectron variant with the highest ON/OFF ratio, we sought to gain deeper insights into its structural determinants and establish generalizable design principles. To achieve this, we leveraged data from the Detectron variant library to develop a machine learning model capable of predicting ON/OFF ratios from Detectron ncRNA structures. For machine learning, we utilized the Extreme Gradient Boosting (XGBoost) algorithm which builds an ensemble of decision trees sequentially, with each new tree correcting the errors made by the previous one^23^. To evaluate model performance and identify the optimal feature set, we employed 5-fold nested cross-validation, reserving 20% of the data in each outer fold for testing and using the remaining 80% for training and inner-loop hyperparameter tuning (**Fig. 3a, Extended Data Fig. 5a**). Performance metrics were averaged across the 5 outer folds to provide an unbiased estimate of generalization. A model trained with the basic five features (stem1 length, stem2 length, bulge length, loop length, d length) exhibited strong performance and prediction accuracy, with an average coefficient of determination (R^2^=0.773) and Pearson correlation coefficient (r=0.880). Adding total stem length (stem1 + stem2 length) as well as the free energy of the hairpin (ΔG_hairpin_) yielded a statistically significant improvements in model performance and prediction accuracy, achieving a coefficient of determination (R^2^=0.794) and Pearson correlation coefficient (r=0.891) (**Fig. 3a,b and Extended Data Fig. 5b-d**).

**Figure 3.**
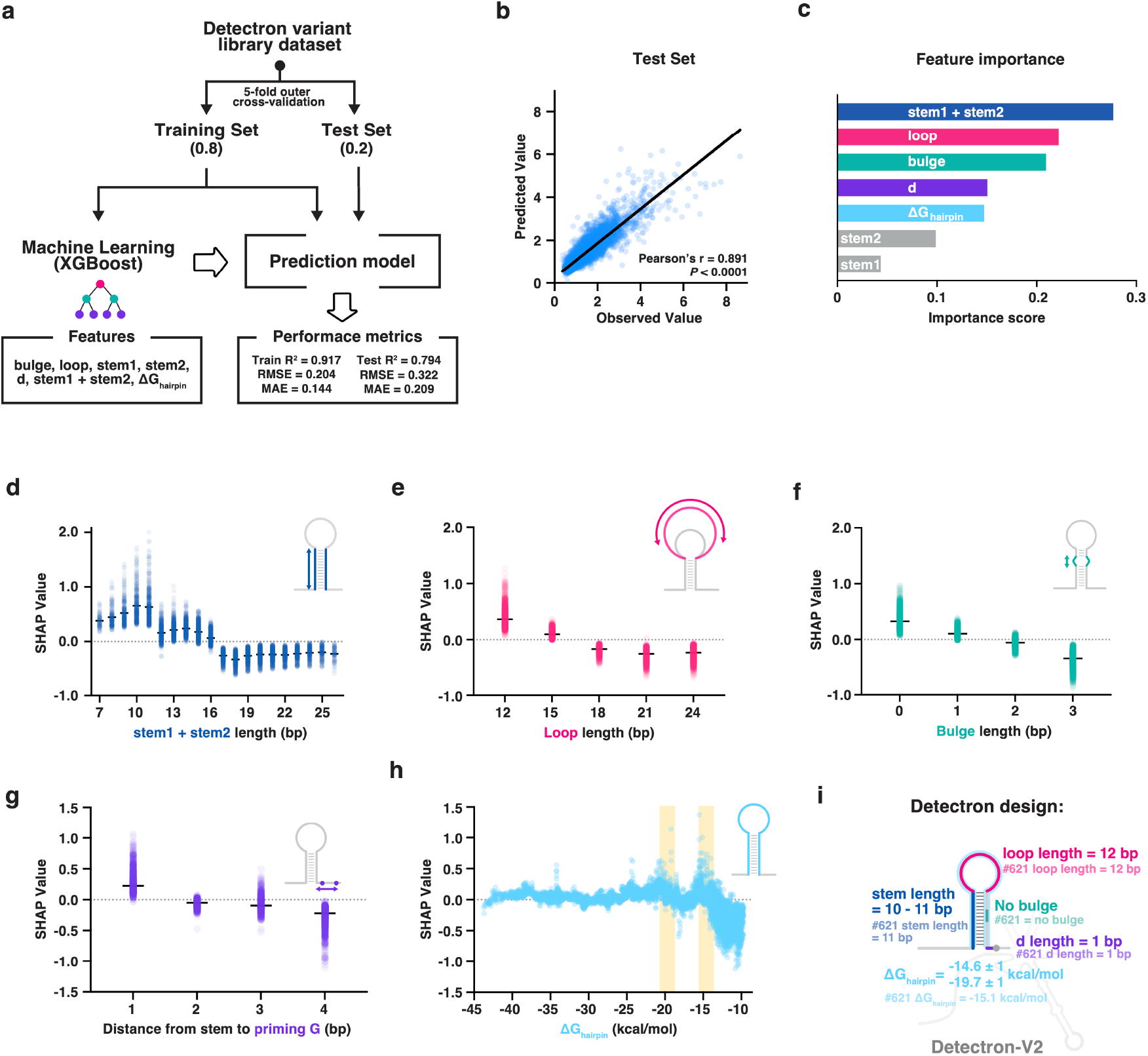
Machine learning on a Detectron variant library identifies key design principles for optimal ON/OFF ratios. **a**, Schematic of the machine learning pipeline using XGBoost, trained on a feature set comprising stem1, stem2, bulge, loop, d, stem1 + stem2, ΔG_hairpin_. Model performance is evaluated using coefficient of determination (R^2^), root mean squared error (RMSE), and mean absolute error (MAE). XGBoost, extreme gradient boost. **b**, Predicted versus observed ON/OFF ratios for the training (left) and test (right) sets. Each circle represents an individual Detectron variant. Pearson’s r and associated *P*-values are annotated on the plot. **c**, Feature importance plot ranking the contribution of each input variable to model predictions. **d-g**, SHAP analysis of ON/OFF ratios across different feature distributions: **d**, combined stem lengths; **e**, loop lengths; **f**, bulge lengths; and **g**, distance from stem to priming G. **h**, free energy of the hairpin (ΔG_hairpin_). Each circle represents an individual Detectron variant. Bars represent the mean SHAP value of the variants. **i**, Extracted design principles for optimal ON/OFF ratios.

For interpretability, we retrained the 7-feature model on all 7,814 variants using the hyperparameters selected during cross-validation. We then applied Shapley Additive Explanations (SHAP) to quantify the contribution of each feature to model predictions across the full dataset, providing the most stable and representative feature importance estimates for guiding future Detectron design. The feature importance analysis revealed that the total stem length was the most influential factor in predicting the ON/OFF ratio of Detectron variants (importance score = 0.277) (**Fig. 3c**). Loop length (0.222) and bulge length (0.209) followed closely, underscoring that ON/OFF switching is governed by multiple structural elements rather than a single dominant feature (**Fig. 3c**). The distance from the stem to the priming guanosine (0.151) and ΔG_hairpin_ (0.147) also contributed meaningfully, suggesting that both spatial configuration and overall hairpin stability influence Detectron performance (**Fig. 3c**). In contrast, stem2 (0.099) and stem1 (0.043) had relatively lower importance scores, likely because their individual contributions were captured by the total stem length feature (**Fig. 3c** and **Extended Data Fig. 6a-b**). These results indicate that the ON/OFF switching mechanism is shaped by a synergistic interplay between stem architecture, loop and bulge size, and the overall thermodynamic stability of the ncRNA structure, rather than being dictated by any single structural element.

Finally, to determine the optimal feature ranges for maximizing ON/OFF ratios, we analyzed SHAP values and the average ON/OFF ratios across different feature values. The highest SHAP values and the average ON/OFF ratios were observed when (i) the combined stem length was 10–11 bp, (ii) the bulge length was 0 bp, (iii) the loop length was 12 bp, and (iv) the d length was 1 bp (**Fig. 3d-g** and **Extended Data Fig. 6c-f**). Among the top 100 variants with the highest SHAP values and average ON/OFF ratios, two dominant peaks in ΔG_hairpin_ were observed at -14.6 ± 1 kcal/mol and -19.7 ± 1 kcal/mol, indicating the presence of two distinct thermodynamic optima that favor efficient switching (**Fig. 3h** and **Extended Data Fig. 6g**). Collectively, these findings demonstrate that the identified structural parameters not only contribute to the model’s prediction of higher ON/OFF ratios but also correlate with experimentally observed enhancements in ON/OFF performance. Notably, the structure of our top performing variant in the library, #621 aligned with all these optimal structural parameters (**Fig. 3i**). Therefore, we used these parameters for our optimized Detectron (Detectron-V2) and adopted its design scheme for subsequent experiments.

### Programmable Detectron-V2 achieves enhanced ON/OFF ratios with high specificity across phage transcripts

Having optimized Detectron design principles using a high-throughput library and identified structural features that strongly influence ON/OFF switching, we next sought to test whether these design rules were generalizable across distinct biological targets. Up to this point, the Detectron performance had been evaluated using a synthetic Zika trigger RNA fragment (135 nt containing the 33-nt target site). To assess programmability and real-world applicability, we applied the optimized Detectron-V2 scheme to design new devices targeting phage transcripts (*gp23* from T4, *gp8* from T5, and *gp10A* from T7) and evaluated using their cognate full-length mRNAs (**Fig. 4a**).

**Figure 4.**
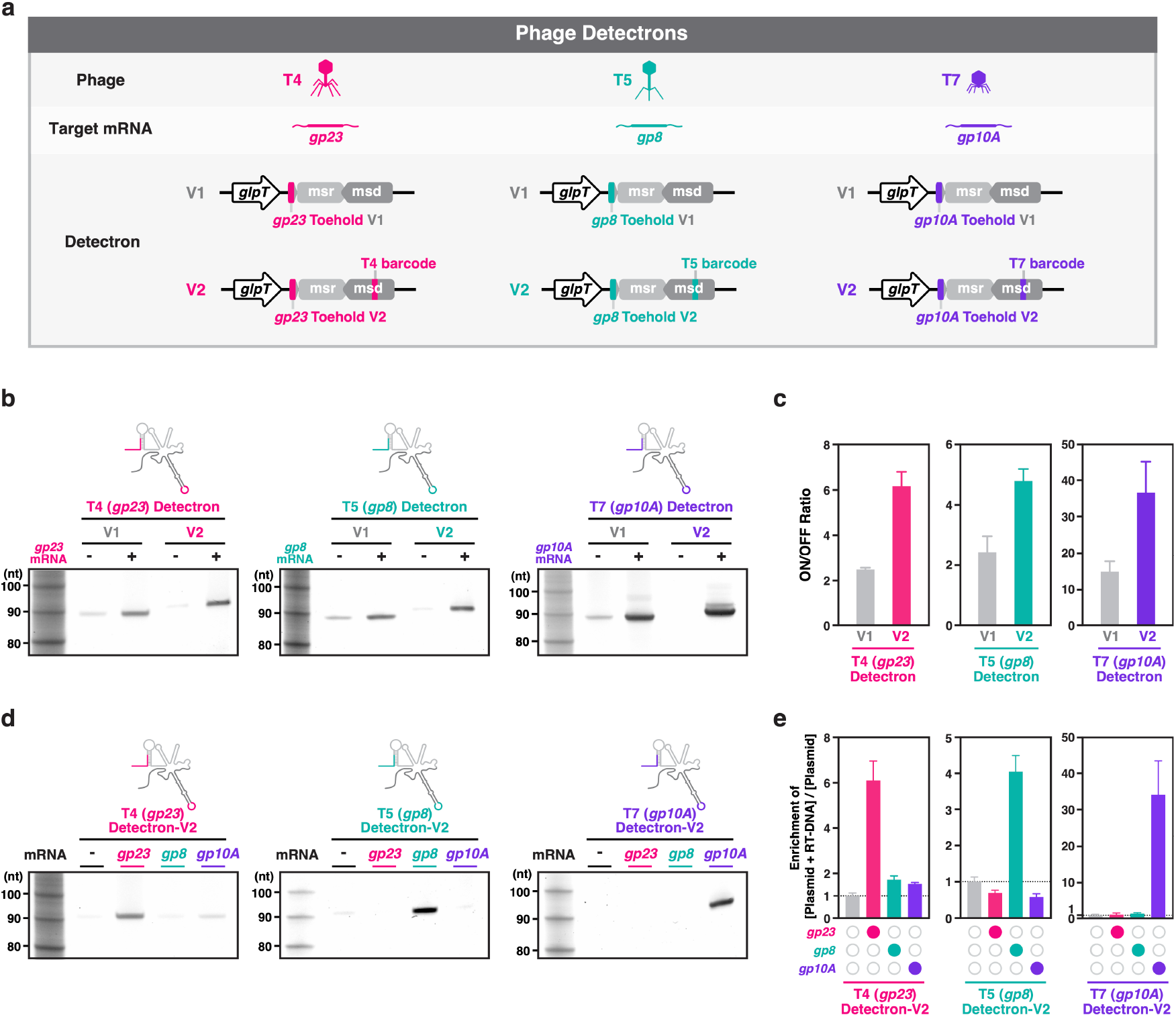
Programmable Detectron-V2 achieves enhanced ON/OFF ratios with high specificity across phage transcripts. **a**, Schematic of Detectrons targeting T4 (*gp23*), T5 (*gp8*), and T7 (*gp10A*) transcripts, showing original (V1) and optimized (V2) constructs. **b**, Urea PAGE analysis of RT-DNA from the indicated Detectrons in the absence (lane 1, 3) or presence (lane 2, 4) of the cognate mRNA expressed under the J23100 promoter. **c**, Quantification of ON/OFF ratios of indicated Detectrons as measured by qPCR; Student’s *t*-test with Welch’s correction (unpaired, two-tailed): T4 Detectron-V2 versus -V1: *P* = 0.009; T5 Detectron-V2 versus -V1: *P* = 0.0045; T7 Detectron-V2 versus -V1: *P* = 0.037. **d**, Urea PAGE analysis of RT-DNA from the indicated Detectrons in the absence (lane 1) or presence of *gp23, gp8*, and *gp10A* mRNA expressed under the J23100 promoter (lane 2, 3, and 4 respectively). **e**, Enrichment of RT-DNA + plasmid over plasmid alone, as measured by qPCR; One-way ANOVA with Tukey’s multiple comparisons test (corrected): T4 Detectron-V2 + *gp23* versus T4 Detectron-V2: *P* < 0.0001; T4 Detectron-V2 + *gp23* versus T4 Detectron-V2 + *gp8*: *P* < 0.0001; T4 Detectron-V2 + *gp23* versus T4 Detectron-V2 + *gp10A*: *P* < 0.0001; T5 Detectron-V2 + *gp8* versus T5 Detectron-V2: *P* < 0.0001; T5 Detectron-V2 + *gp8* versus T5 Detectron-V2 + *gp23*: *P* < 0.0001; T5 Detectron-V2 + *gp8* versus T5 Detectron-V2 + *gp10A*: *P* < 0.0001; T7 Detectron-V2 + *gp10A* versus T7 Detectron-V2: *P* = 0.0001; T7 Detectron-V2 + *gp10A* versus T7 Detectron-V2 + *gp23*: *P* = 0.0001; T7 Detectron-V2 + *gp10A* versus T7 Detectron-V2 + *gp8*: *P* = 0.0001. Bars represent the mean (±s.d.) of three biological replicates.

We compared the performance of these phage-targeting Detectrons with their original V1 counterparts. In all cases, Detectron-V2 exhibited a clear reduction in OFF-state signal and an increase in ON-state output, leading to significantly enhanced ON/OFF ratios relative to V1 (**Fig. 4b,c**). These results confirm that the structural principles defined in the Detectron variant library generalize robustly, as exemplified here by phage transcripts.

To further evaluate specificity, we challenged each Detectron-V2 with both cognate and non-cognate phage transcripts. Each device was strongly activated only by its cognate phage transcript with negligible responses to non-cognate inputs, demonstrating high specificity and low cross-activation across the T4, T5, and T7 Detectrons (**Fig. 4d-e**).

### Detectrons enable discrimination of phage infections in a single-pot culture

To evaluate whether Detectrons can distinguish phage infections in a single-pot culture, we utilized the three phage-targeting Detectrons each with a unique 8-nt DNA barcode encoded into the msd loop (**Fig. 5a**). In parallel, we included an *E. coli* strain harboring a No Toehold retron-Eco1, which constitutively produces a wt loop barcode. This strain functions as both a positive control and a normalization reference for sequencing-based quantification.

**Figure 5.**
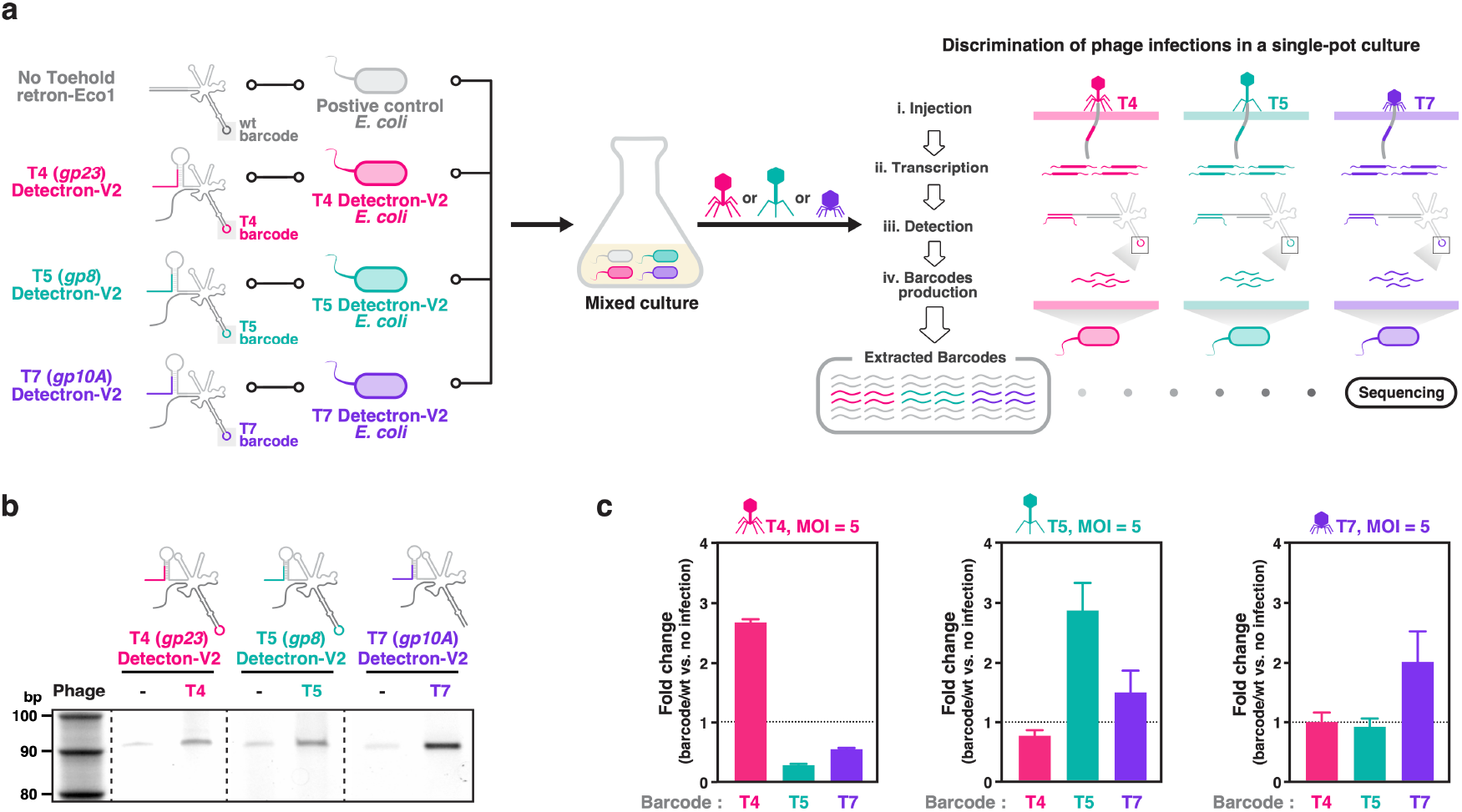
Detectrons enable discrimination of phage infections in a single-pot culture. **a**, Schematic overview of the sequencing-based phage detection workflow. A mixed *E. coli* culture containing strains expressing phage-specific Detectrons—targeting T4 (*gp23*), T5 (*gp8*), and T7 (*gp10A*)—or a No Toehold retron-Eco1 control was challenged with either T4, T5, or T7 phages. Following infection, intracellular RT-DNAs were isolated and analyzed by high-throughput sequencing. **b**, Urea PAGE analysis of RT-DNA from the indicated Detectron strains in the absence (lane 1, 3, 5) or presence (lane 2, 4, 6) of cognate phage infection. Chloramphenicol was added post-infection to prevent premature lysis—at 10 min for T4 and T7, and 25 min for T5—to enable intracellular accumulation of Detectron-derived RT-DNA. **c**, Quantification of barcode induction by sequencing. Fold-change in barcode abundance relative to the no-infection control is shown for each phage challenge; One-way ANOVA with Tukey’s multiple comparisons test (corrected): [T4 infection] T4 barcode versus T5 barcode: *P* < 0.0001; T4 barcode versus T7 barcode: *P* < 0.0001; [T5 infection] T5 barcode versus T4 barcode: *P* = 0.0003; T5 barcode versus T7 barcode: *P* = 0.0023; [T7 infection] T7 barcode versus T4 barcode: *P* = 0.0189; T7 barcode versus T5 barcode *P* = 0.0134; Bars represent the mean (±s.d.) of three biological replicates.

We hypothesized that barcode sequencing from a pooled *E. coli* culture—comprising strains bearing phage-specific Detectrons or the No Toehold retron control—after phage challenge would enable identification of the infecting phage species (**Fig. 5a**). Each Detectron carrying strain was first challenged with its cognate phage to evaluate whether Detectrons can detect phage-derived mRNA expressed during infection, representing a shift from synthetic to physiologically relevant RNA detection (**Extended Data Fig. 7a**).

However, detecting phage infection poses a timing challenge: sufficient time is needed post-infection for DNA barcodes to accumulate, yet prolonged incubation risks extensive cell lysis and barcode loss into the medium (**Extended Data Fig. 7b**). Conversely, early harvest may fail to capture sufficient barcode signal (**Extended Data Fig. 7c**). To resolve this tradeoff, we introduced chloramphenicol post-infection to inhibit bacterial translation. This antibiotic blocks peptidyl transferase activity of the ribosome, which will halt phage protein synthesis and delay lysis. We reasoned that this intervention would provide a temporal window during which Detectron-derived DNA barcodes could accumulate intracellularly without immediate lysis, improving sensitivity and recovery (**Extended Data Fig. 7d**).

With chloramphenicol treatment, the accumulation of DNA barcodes was observed in all three Detectron carrying strains in response to the cognate phage infection, as visualized via PAGE confirming that Detectrons can sense phage-derived mRNAs transcribed during the infection process (**Fig. 5b**).

To assess the system’s capacity for multiplexed detection, we infected the mixed *E. coli* culture with either T4, T5, or T7 phages. Following infection, DNA barcodes were extracted and quantified to determine which Detectrons had been activated (**Fig. 5a**). In each case, barcode signals corresponding only to the infecting phage were detected, with negligible off-target activation (**Fig. 5c**). This demonstrates that Detectrons can accurately distinguish phage infections within a single culture vessel through barcode-based readouts.

### Detectrons enable multiplexed assessment of host susceptibility to phage infection through dual DNA barcoding

Finally, to evaluate whether Detectrons can quantify host susceptibility to phage infection, we implemented a dual barcoding strategy that simultaneously encoded (1) phage identity and (2) host strain identity within the msd loop of the Detectron, focusing on phage T7 to evaluate host susceptibility across strains (**Fig. 6a**). We used multiple *E. coli* strains with varying disruptions to the lipopolysaccharide (LPS) biosynthesis pathway, which serves as a primary reversible receptor for T7 phage adsorption. As LPS truncations impair phage infectivity to different extents depending on the region disrupted, this system provides a model to test infection sensitivity across strains. Specifically, Δ*rfaG* lacks the outer core of LPS and should moderately inhibit T7 infection, while Δ*rfaD* results in severe truncations affecting the inner core that should strongly inhibit T7 infection (**Fig. 6b**).

**Figure 6.**
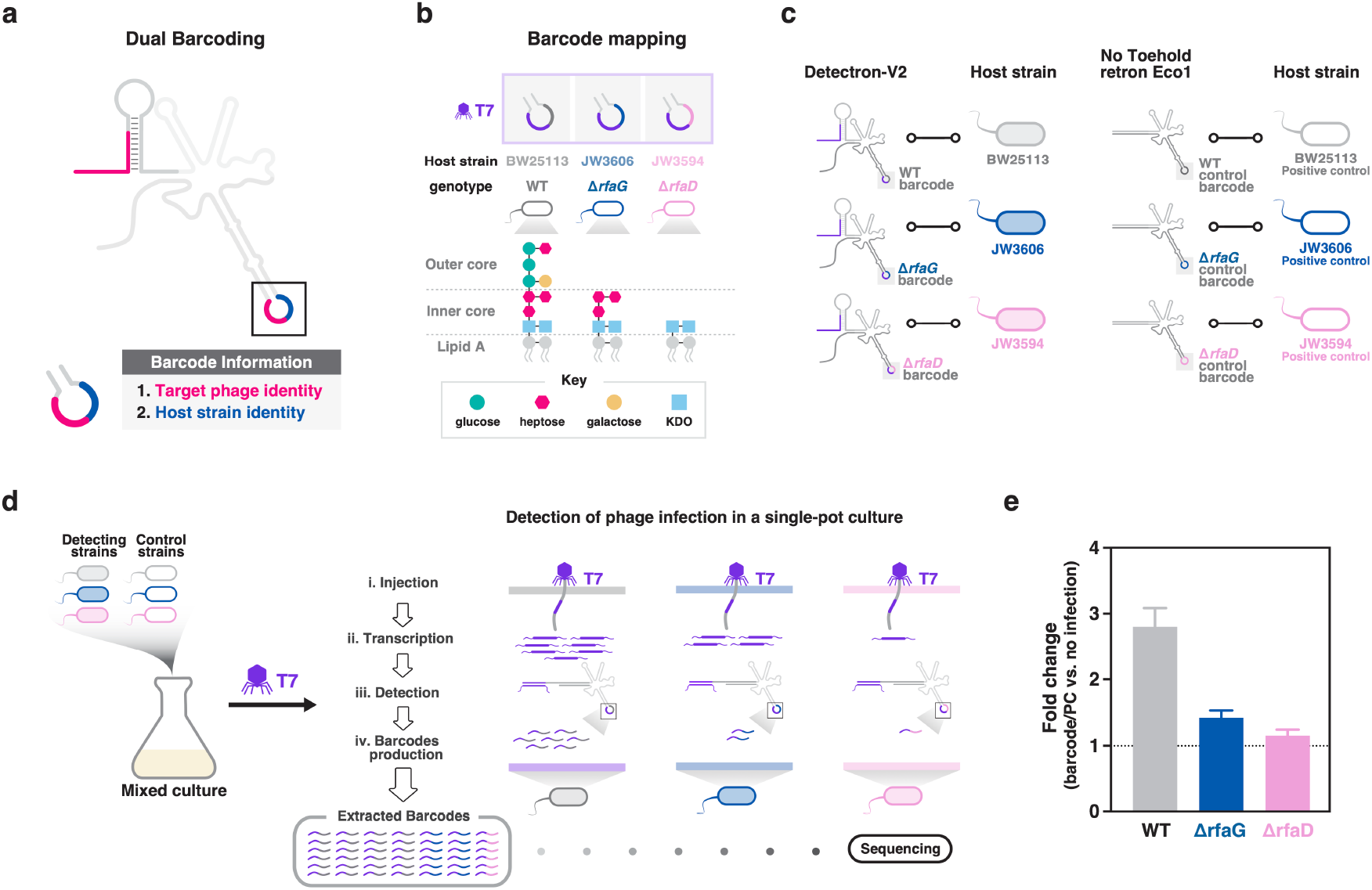
Detectrons enable multiplexed assessment of host susceptibility to phage infection through dual DNA barcoding. **a**, Schematic overview of dual barcoding strategy. Two pieces of information are encoded in the loop of msd; (1) the identity of the target phage and (2) the identity of the host strain. **b**, Barcode mapping of the three host trains. LPS core structures are shown for each strain, highlighting differences in outer core sugars resulting from mutations in Δ*rfaG* and Δ*rfaD*. **c**, Experimental design of the multiplexed phage infection assay. Three T7-targeting Detectron-V2 devices, each encoding a strain-specific barcode, were introduced into wt, Δ*rfaG*, and Δ*rfaD* strains, respectively. In parallel, No Toehold retron-Eco1 constructs with matching strain-specific barcodes served as infection-independent positive controls for normalization. **d**, Detectron-based detection of phage infection in a single-pot culture. A pooled mixture of detecting and control strains was challenged with T7 phage. Upon infection, phage mRNA activates the corresponding Detectrons, triggering barcode production. Following infection, barcodes were extracted and sequenced to quantify infection-specific signal for each host strain. **e**, Quantification of infection susceptibility across strains. Fold-change in barcode abundance relative to the no-infection control is shown; One-way ANOVA with Tukey’s multiple comparisons test (corrected): wt barcode versus Δ*rfaG* barcode: *P* = 0.0002; wt barcode versus Δ*rfaD* barcode: *P* < 0.0001; Bars represent the mean (±s.d.) of three biological replicates.

We engineered three T7-targeting Detectrons, each encoded with a strain-specific DNA barcode, and introduced them into *E. coli* wt (BW25113), Δ*rfaG* (JW3606), and Δ*rfaD* (JW3594) strains, respectively (**Fig. 6c**). In parallel, we included three No Toehold retron-Eco1 controls, each with a corresponding strain-specific barcode, and introduced them into the same strains. These served as both positive controls and normalization references for sequencing-based quantification. The mixed culture containing all six strains was then challenged with T7 phage. Following infection, DNA barcodes were extracted and sequenced to determine the degree of activation of each Detectron (**Fig. 6d**). Barcode signal intensity correlated with expected infectivity: the wt strain exhibited the strongest barcode induction, while Δ*rfaG* and Δ*rfaD* strains showed progressively diminished responses (**Fig. 6e**). This gradient aligned with the known impact of LPS truncation onT7 adsorption efficiency. Together, these results demonstrate that Detectrons not only detect phage infection with high specificity but also enable quantitative assessment of infection susceptibility across genetically distinct bacterial populations, all within a single assay.

## DISCUSSION

The Detectron technology introduces a fundamentally distinct biosensing paradigm by coupling RNA recognition to retron-mediated DNA barcoding. Unlike previous toehold switches, which translate RNA inputs into protein-based readouts such as fluorescence reporters^9^, or CRISPR-based detection systems, which rely on programmable nucleic acid cleavage and collateral reporter activation^24–26^, or those that use engineered crRNAs or guide RNAs to indirectly sense RNA expression via DNA targets^27^, Detectrons directly generate a DNA signal. This retron-derived DNA barcode can be readily quantified by sequencing, enabling scalable and multiplexed detection in pooled setting. Such a DNA-based output eliminates reliance on protein or fluorescence-based reporters and provides a stable molecular signal, making Detectrons particularly suited for high-throughput applications where digital readout is required. Moreover, this DNA output could, in principle, be adapted into molecular recording frameworks, extending Detectron’s utility beyond transient detection.

Our high-throughput variant library analysis established design principles for maximizing ON/OFF ratios. We found that combined stem length, bulge length, loop length, and hairpin stability jointly govern Detectron performance. Incorporating these principles into an optimized architecture (Detectron-V2) substantially improved ON/OFF switching compared to the original design (V1). Importantly, these design rules generalized robustly: variant #621, which exemplifies the optimized Detectron-V2 design, conferred enhanced ON/OFF ratios not only in *E. coli*, but also in the clinically relevant strains *K. pneumoniae* and *C. freundii*. When applied to build phage-targeting Detectrons, the V2 architecture consistently outperformed V1 across three distinct mRNA targets (*gp23, gp8, gp10A*), demonstrating that the design principles defined in the Detectron variant library are both portable across species and programmable across RNA targets.

Building upon these improvements, Detectron-V2 also exhibited high target specificity. Each phage-targeting device responded selectively to its cognate mRNA while exhibiting negligible cross-activation by non-cognate inputs. Leveraging this specificity, we enabled multiplexed detection of specific phage infections and quantitative assessment of host susceptibility within a pooled bacterial culture. These results highlight the potential for Detectrons as a versatile tool for multiplexed RNA biosensing applications where precise discrimination among diverse RNA signals is critical.

To our knowledge, Detectrons represent the first synthetic biosensor that directly converts RNA inputs into DNA barcodes through retron-mediated reverse transcription. Another potential use of this DNA could come in molecular recorders, which have relied on promoter-driven expression to generate the input to the ‘write’ operation, using tools such as CRISPR-Cas-based genome editing^28–31^, base editors^32,33^, integrases^34–36^, or site-specific recombinases^37,38^—to encode biological events indirectly. Detectrons bypass this limitation by enabling programmable RNA sensing and immediate DNA barcode synthesis upon target transcript detection. This direct RNA-to-DNA conversion introduces a new modality for RNA-responsive devices: transient RNA states can now be captured as durable, sequence-encoded DNA records, readily retrievable by sequencing.

While our study focuses on bacterial systems, retrons have recently been shown to function in mammalian cells as DNA donors for genome editing^15–20^. This raises the exciting possibility that Detectrons could be adapted to eukaryotic contexts to detect endogenous transcripts and permanently record them as DNA barcodes. Such a system could enable in vivo transcriptional surveillance, lineage tracing, and diagnostic monitoring in higher organisms. More broadly, the combination of programmable RNA sensing with retron-based DNA recording expands the synthetic biology toolkit beyond protein or RNA outputs, introducing a fundamentally new axis of information transfer that bridges RNA dynamics with stable DNA memory.

## METHODS

All biological replicates were taken from distinct samples, not the same sample measured repeatedly. For Detectron variant libraries, each biological replicate is an independent electroporation and expression of the libraries into the strain *E. cloni* 10G. All statistical tests and *P*-values are included in Supplementary Table S1.

### Constructs and strains

Bacterial experiments were performed in bMS.346 which was generated from *E. coli* MG1655 by inactivating the *exoI* and *recJ* genes with early stop codons as described in previous work^39^. *Klebsiella pneumoniae* (ATCC 10031) and *Citrobacter freundii* (ATCC 8090) were used to assess portability of Detectron-V2. Bacteriophages T4 (ATCC 11303-B4), T5 (ATCC 11303-B5), and T7 (ATCC BAA-1025-B2) were used to evaluate the capability of Detectrons to detect phage-derived mRNAs in infected *E. coli* cells.

### RT-DNA purification and PAGE analysis

To analyze RT-DNA on a PAGE gel, 20 ml of culture was pelleted, and RT-DNAs were extracted using the QIAGEN Midiprep Plasmid Plus Kit. The purified RT-DNAs were then analyzed on 10% Novex TBE-Urea gels (Invitrogen) with 1× TBE running buffer that was heated to >80°C before loading. Gels were stained with SYBR Gold (Thermo Fisher) and imaged on a Gel Doc imager (Bio-Rad).

### qPCR

qPCR analysis of RT-DNA was performed using two primer sets: (1) Eco1_wt_2F and Eco1_wt_R, and (2) Eco1_wt_3F and Eco1_wt_R. The first set amplifies both plasmid DNA and RT-DNA, as the primers bind within the msd region, while the second set selectively amplifies plasmid DNA by targeting a region outside the msd. The results were analyzed by first taking the difference in cycle threshold (Ct) between the two primer sets for each biological replicate. The relative enrichment of [Plasmid + RT-DNA] compared to [Plasmid] within each sample was calculated as 2^-ΔCt^.

### Variant library cloning

The synthesized oligos were first amplified using skpp15 primer sets. The Detectron variant library was constructed using a two-step Golden Gate cloning strategy: in the first step, the amplified oligos were cloned into the backbone using SapI Type IIS restriction enzyme; in the second step, the invariable region—comprising the msr and the first 29 bp of msd—was reinserted using BsaI Type IIS restriction enzyme, restoring the complete Detectron construct. Two libraries were constructed using this method: an ON library (constitutively expressing trigger RNA) and an OFF library (no trigger RNA expression). 50 μl of *E. cloni* 10G cells were added to 0.1-cm gap electroporation cuvettes (BioRad #1652083), and 6 μl of cleaned-up Golden Gate reaction was mixed with the cells. Cells were electroporated with the following settings: 1800 V, 25 μF, 200 ν. was electroporated with these libraries for RT-DNA production. After the pulse, cells were quickly recovered in 1 mL of pre-warmed recovery medium and placed in a shaking incubator for 1 h at 37°C. Dilutions of the recovered cells were plated on LB + carbenicillin plates to estimate CFUs, and the rest of recovered cells were transferred into 24 mL of pre-warmed LB + carbenicillin and grown for overnight. The next day, CFUs were estimated, and the experiments were continued only if the library coverage was estimated to be >200×.

### Variant library expression and sequencing

After an overnight culture, barcode-encoded RT-DNAs were extracted using the QIAGEN Midiprep Plasmid Plus Kit. From the Midiprep eluate, 10 μl was reserved for plasmid quantification, while the remaining 190 μl was purified with the Zymo ssDNA Clean & Concentrator Kit to deplete plasmid DNA. Using a previously described sequencing pipeline to quantify RT-DNA abundance^16–18,22^, two rounds of PCR amplification were performed to add sequencing adapters and indexes to the extracted RT-DNAs and plasmids, and the products were sequenced using an Illumina platform (NextSeq 2000). Illumina FASTQ files were demultiplexed and processed using custom code (https://github.com/Shipman-Lab/Detectron). The abundance of RT-DNAs from each variant was normalized to the abundance of its corresponding plasmid, which were prepped and sequenced in parallel.

### Machine learning analysis of Detectron variant performance

Data from the Detectron variant library were analyzed to develop a predictive machine learning model. Four feature sets were tested: (1) a 5-feature set (stem1 length, stem2 length, bulge length, loop length, d length), (2) a 6-feature set A (5 features + total stem length (stem1 + stem2)), (3) a 6-feature set B (5 features + hairpin free energy (ΔG_hairpin_)), and (4) a 7-feature set (all features combined). For each model, features were standardized with StandardScaler fit only on the training split within each fold and then applied to the corresponding validation/test split.

Models were trained using XGBoost regression (XGBRegressor, v1.6.2) and hyperparameters were selected with nested 5-fold cross-validation: an outer 5-fold split provided held-out test folds, and within each outer training split an inner 5-fold GridSearchCV was used to tune hyperparameters. Model performance was assessed on the outer test folds using five key metrics: R^2^, RMSE, MAE, Pearson correlation coefficient r, and *P*-value (statistical significance of correlation). Mean performance values across folds were calculated for each feature set, and paired *t*-tests were performed to determine whether differences in model performance were statistically significant (*P* < 0.05 was considered significant).

After identifying the best-performing feature set from the cross-validated results, a final model was retrained on all samples using the hyperparameters selected by the inner CV. To interpret the model’s predictions, Shapley Additive Explanations (SHAP) analysis was conducted using shap.TreeExplainer. Feature contributions were ranked based on mean absolute SHAP values, and SHAP summary plots and feature importance rankings were generated to visualize the effect of individual features on ON/OFF ratio prediction.

### Phage target mRNA sequence selection

Phage-derived mRNA target sequences were identified using a customized version of the Toehold Designer tool (https://github.com/EPFLliGem/Toehold-Designer). The secondary structure template of the toehold switch was redesigned to match the structure of Detectron-V2. In addition, full-length phage mRNAs were segmented using a sliding window of 33 nucleotides—rather than the default 36-nt window—to align the target length with the structural parameters of the Detectron-V2.

### Detection of phage infections in a single-pot culture

Four *E. coli* strains were generated, each expressing either the No Toehold retron-Eco1 (producing a wt barcode) or a phage-specific Detectron-V2 targeting T4 (*gp23*), T5 (*gp8*), or T7 (*gp10A*) mRNA. At OD600 = 0.4, the strains were mixed at equal ratios and infected with either T4, T5, or T7 phage at a multiplicity of infection (MOI) of 5. Cultures were incubated at 37°C with shaking for 1 hour.

To enhance intracellular barcode retention, chloramphenicol was added post-infection to inhibit bacterial translation and delay lysis: 25 minutes post-infection for T5, and 10 minutes post-infection for T4 and T7. After 1 hour, cultures were centrifuged at 4,000 rpm for 20 minutes at 4°C. Barcode-encoded RT-DNAs and plasmids were extracted as described above and sequenced using either the Illumina NextSeq 2000 or Oxford Nanopore PromethiON platforms. FASTQ files were demultiplexed and processed using custom codes (https://github.com/Shipman-Lab/Detectron). The abundance of barcode-encoded RT-DNAs were normalized to the abundance of the corresponding plasmid.

For barcode quantification, phage-specific barcode read counts were first normalized to wt barcode reads using the formula:

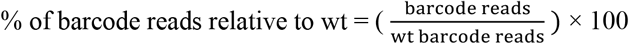

Subsequently, fold change was calculated by comparing infected versus non-infected samples:

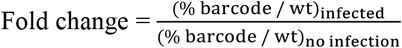

### Multiplexed assessment of host susceptibility to phage infection

Three T7-targeting Detectrons, each harboring a strain-specific DNA barcode, were engineered and introduced into wt, Δ*rfaG*, and Δ*rfaD E. coli* strains, respectively. Three No Toehold retron-Eco1 controls, each with a corresponding strain-specific barcode were introduced into the same strains. When cultures reached an OD600 of 0.4, detecting and control strains were mixed in a 4:4:4:1:1:1 ratio (detecting:control = 4:1) and infected with T7 phage at a multiplicity of infection (MOI) of 5. Cultures were incubated at 37°C with shaking for 1 hour. To enhance intracellular barcode retention, chloramphenicol was added 10 minutes post-infection to inhibit bacterial translation and delay lysis. After 1 hour, cultures were centrifuged at 4,000 rpm for 20 minutes at 4°C. Barcode-encoded RT-DNAs and plasmids were extracted and sequenced as described above.

## ACKNOWLEDGEMENTS

Work was supported by funding from the Bachrach Family Foundation and the Robert J Kleberg, Jr. and Helen C. Kleberg Foundation. S.L.S. is a San Francisco Biohub Investigator. We thank Katherine S. Pollard for discussions regarding machine learning, Kate D. Crawford for discussions regarding Detectron library construction, and Alejandro G. Delgado for assistance in assessing Detectron portability.

## AUTHOR CONTRIBUTIONS

Author contributions follow the CRedIT taxonomy (https://www.elsevier.com/researcher/author/policies-and-guidelines/credit-author-statement)

J.H.: Conceptualization, Methodology, Investigation, Writing, Visualization, Supervision. S.L.S.: Conceptualization, Methodology, Supervision, Project administration, Funding acquisition.

## DATA AND CODE AVAILABILITY

Sequencing data associated with this study are available in the NCBI SRA (PRJNA1366125) https://www.ncbi.nlm.nih.gov/bioproject/PRJNA1366125

Custom code used to process or analyze data from this study is available at: https://github.com/Shipman-Lab/Detectron

## SUPPLEMENTARY MATERIALS

Extended Data File

Supplementary Table 1-3

Supplementary Table 4

