## Extended Data Files for "Programmable RNA detection generates DNA barcodes for multiplexed phage-host interaction screening"

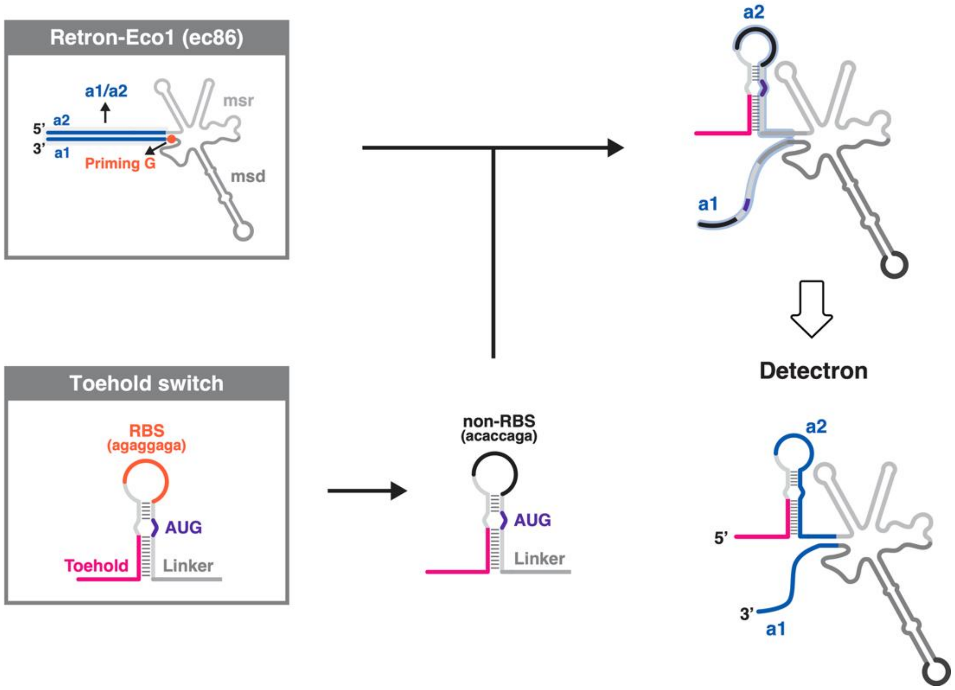


**Extended Data Fig. 1 Structural design of Detectron.** A Zika virus RNA-responsive toehold switch (bottom left) and retron Eco1 (top left) were integrated to construct Detectron. The ribosome-binding site (RBS) within the toehold loop was replaced with a non-RBS sequence to prevent unintended translation of the retron ncRNA. The region spanning from the non-RBS to the linker was repurposed as the a2 domain of the Eco1 ncRNA, with the complementary a1 sequence redesigned accordingly.

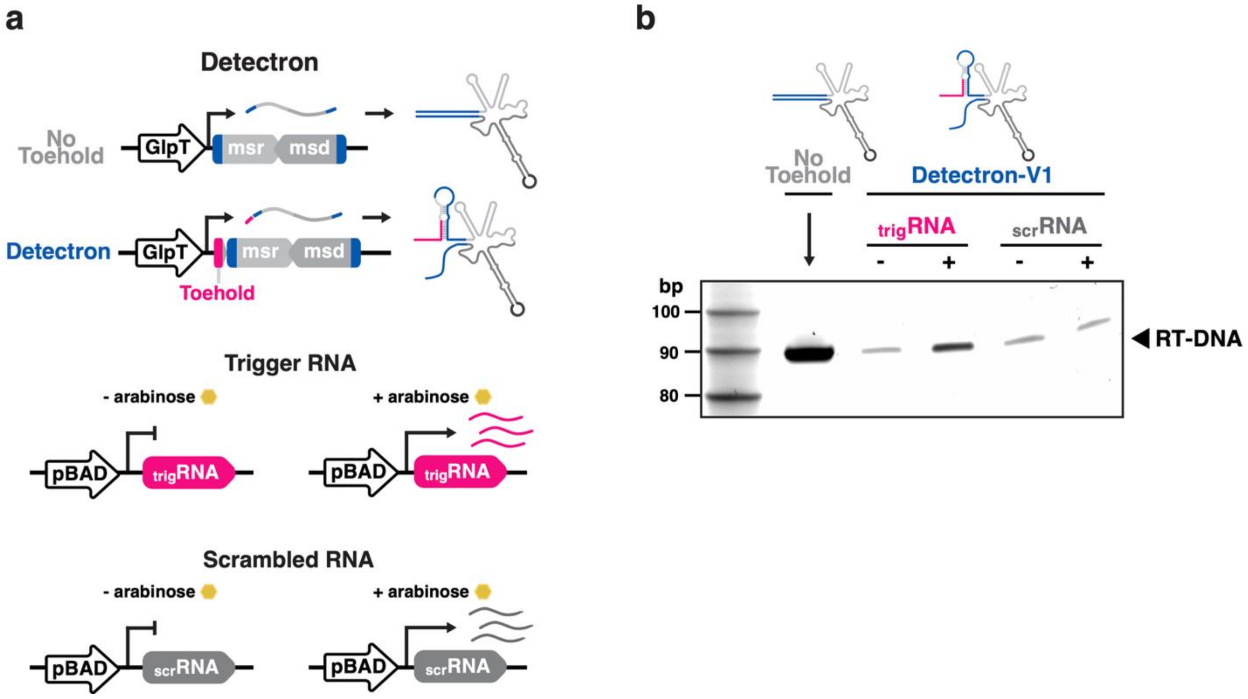


**Extended Data Fig. 2 Detectrons specifically respond to target RNA expressed under an inducible promoter. a**, Schematic of the experimental promoters used to express No Toehold control, Detectron, and trigger RNA, along with representations of the expressed transcripts. msr, multicopy single-stranded RNA. msd, multicopy single-stranded DNA. trigRNA, trigger RNA. **b**, Urea PAGE analysis of RT-DNA from the No Toehold control (lane 1, excluding ladder) and from Detectron in the absence (lane 2) or presence of trigger RNA expressed under the pBAD promoter (lane 3), or in the absence (lane 4) or presence of scrambled RNA expressed under the pBAD promoter (lane5).

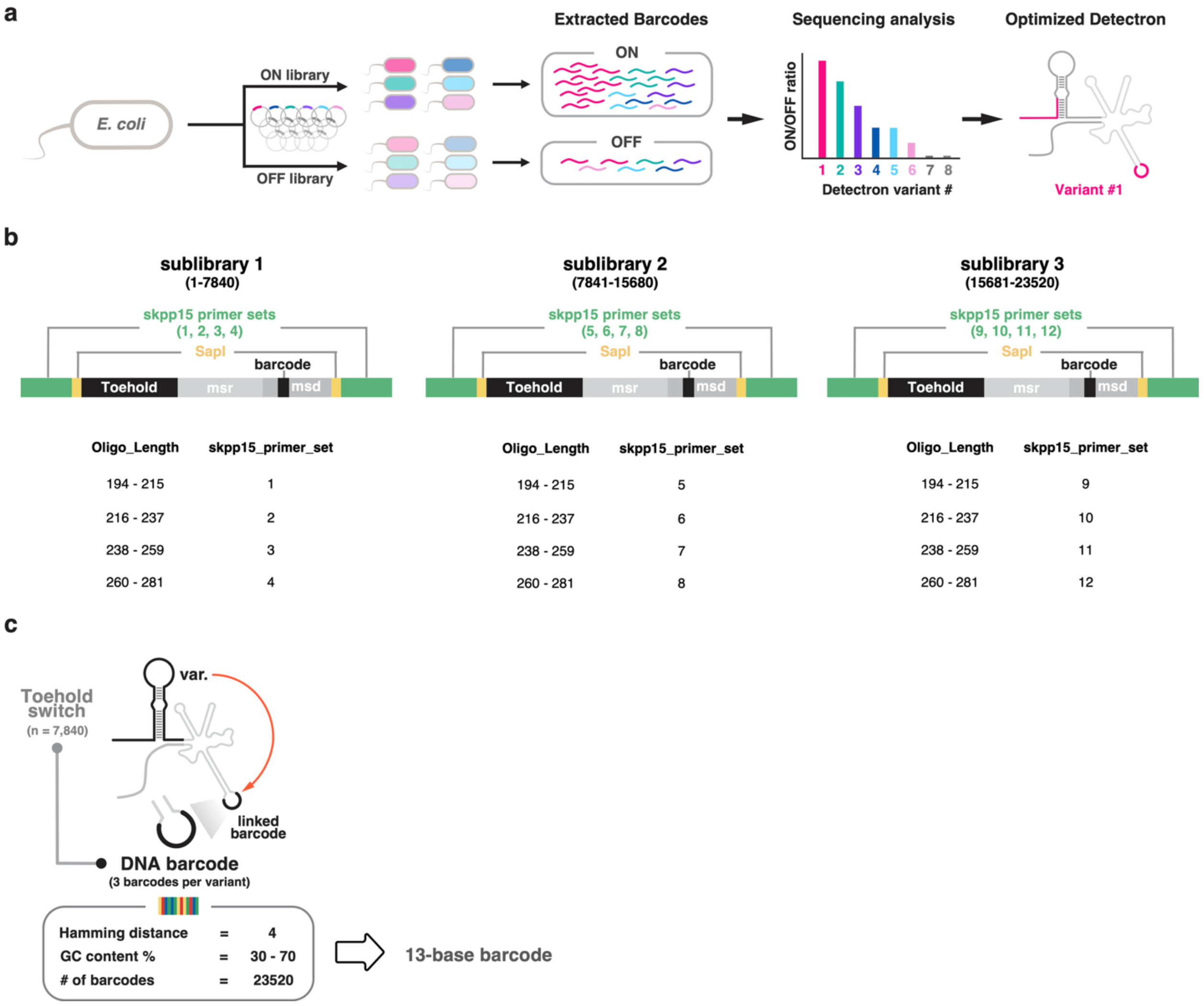


**Extended Data Fig. 3 Design and sequencing pipeline of the Detectron variant library. a**, Schematic overview of the workflow used to identify Detectron variants with the highest ON/OFF ratios. Two libraries were constructed: an ON library (constitutively expressing trigger RNA) and an OFF library (no trigger RNA expression). Both were transformed into *E. cloni* 10G cells for RT-DNA production. Barcode-encoded RT-DNAs were subsequently isolated and sequenced. **b**, Each Detectron variant was linked to three unique DNA barcodes, generating three sublibraries and expanding the total library size to 23,520 distinct constructs. **c**, Design parameters of 13-base DNA barcodes, constrained by a minimum Hamming distance of 4 and GC content between 30% and 70%, to ensure robust sequence differentiation and stability.

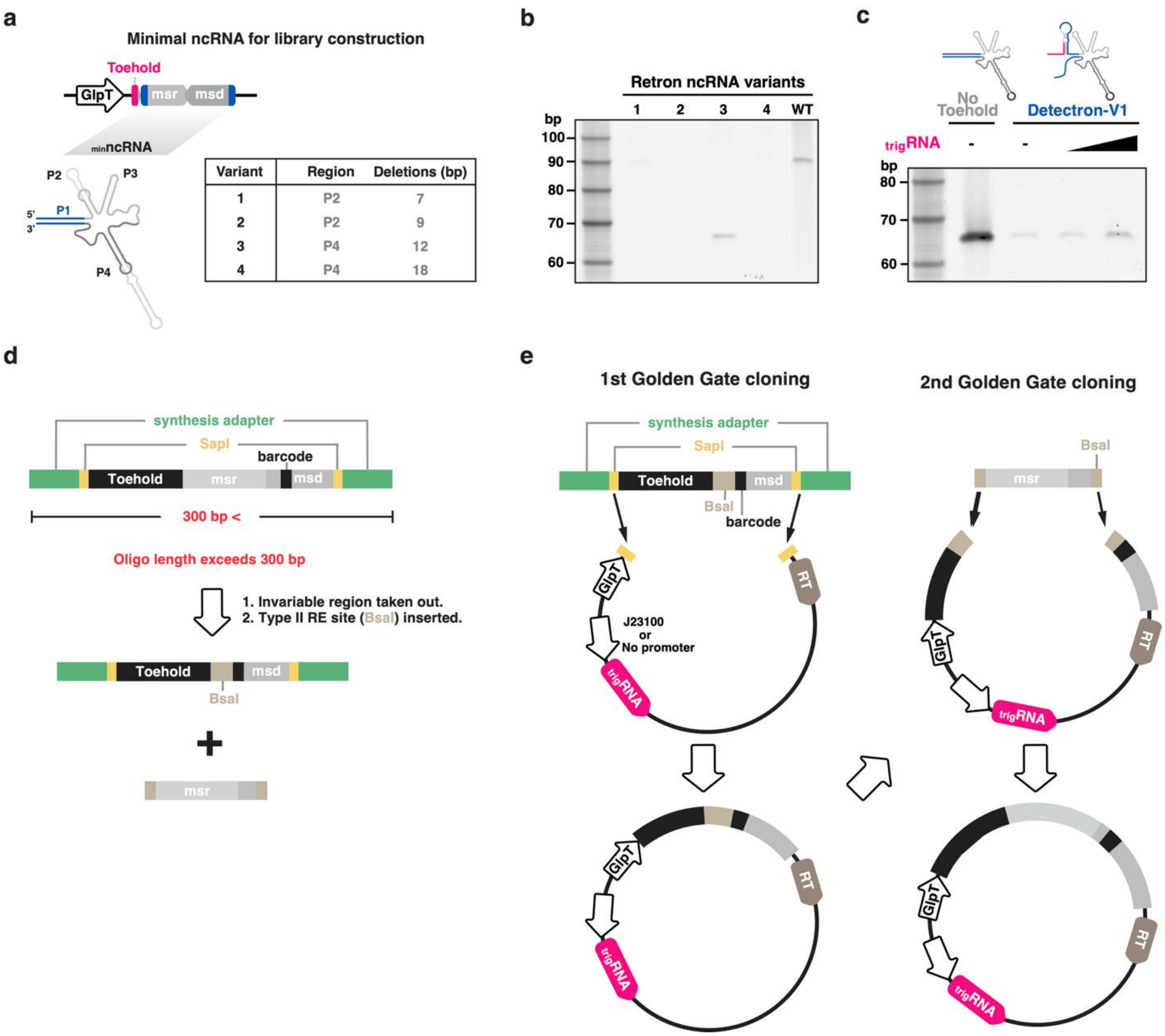


**Extended Data Fig. 4 Construction of the Detectron variant library. a**, Schematic of Retron Eco1 ncRNA variants tested. **b**, Urea PAGE analysis of RT-DNA from the indicated ncRNA variants (lanes 1-4, excluding ladder) and wild-type (WT) ncRNA (lane 5). **c**, Urea PAGE analysis of RT-DNA from the No Toehold control (lane 1, excluding ladder) and Detectron-V1 in the absence (lane 2) or presence of trigger RNA expressed under the J23105 promoter (lane 3) or J23100 promoter (lane 4). **d**, Design of Detectron variant oligonucleotides. The invariable region containing the full msr and partial msd sequences was omitted from synthesized oligos. **e**, Pooled oligos were cloned into either ON (constitutively expressing trigger RNA) or OFF (no trigger RNA expression) backbones via SapI digestion. The invariable region was subsequently reinserted into both ON and OFF libraries via BsaI digestion.

**a**

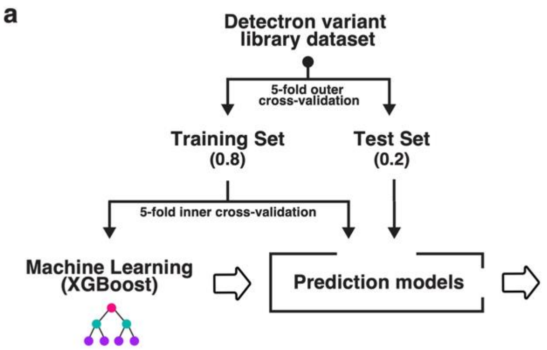


| Feature set | Features inlcuded for learning |
|---|---|
| 5 | bulge, loop, d, stem1, stem2 |
| 6A | bulge, loop, d, stem1, stem2, stem1 + stem2 |
| 6B | bulge, loop, d, stem1, stem2, $\Delta G_{hairpin}$ |
| 7 | bulge, loop, d, stem1, stem2, stem1 + stem2, $\Delta G_{hairpin}$ |

**b**

**Prediction model performance**

| Feature set | Fold set | Model performace metrics (5-fold cross-validation results) | | | | | | | |
|---|---|---|---|---|---|---|---|---|---|
| | | $R^2$ | | RMSE | | MAE | | Pearson r | |
| | | Train | Test | Train | Test | Train | Test | Train | Test |
| 5 | 0 | 0.885 | 0.764 | 0.240 | 0.351 | 0.166 | 0.215 | 0.941 | 0.876 |
| | 1 | 0.879 | 0.795 | 0.246 | 0.326 | 0.167 | 0.218 | 0.938 | 0.892 |
| | 2 | 0.885 | 0.779 | 0.240 | 0.343 | 0.165 | 0.219 | 0.941 | 0.883 |
| | 3 | 0.882 | 0.761 | 0.247 | 0.329 | 0.168 | 0.214 | 0.940 | 0.873 |
| | 4 | 0.889 | 0.767 | 0.238 | 0.339 | 0.164 | 0.225 | 0.943 | 0.876 |
| 6A | 0 | 0.901 | 0.794 | 0.223 | 0.329 | 0.155 | 0.206 | 0.949 | 0.892 |
| | 1 | 0.902 | 0.805 | 0.221 | 0.318 | 0.154 | 0.211 | 0.950 | 0.898 |
| | 2 | 0.901 | 0.799 | 0.222 | 0.327 | 0.155 | 0.210 | 0.950 | 0.894 |
| | 3 | 0.873 | 0.781 | 0.256 | 0.315 | 0.175 | 0.207 | 0.935 | 0.884 |
| | 4 | 0.904 | 0.784 | 0.221 | 0.327 | 0.154 | 0.218 | 0.951 | 0.885 |
| 6B | 0 | 0.915 | 0.780 | 0.206 | 0.339 | 0.147 | 0.213 | 0.957 | 0.884 |
| | 1 | 0.916 | 0.801 | 0.205 | 0.321 | 0.145 | 0.212 | 0.958 | 0.895 |
| | 2 | 0.914 | 0.798 | 0.206 | 0.328 | 0.147 | 0.212 | 0.957 | 0.894 |
| | 3 | 0.915 | 0.787 | 0.209 | 0.311 | 0.148 | 0.205 | 0.957 | 0.887 |
| | 4 | 0.917 | 0.779 | 0.205 | 0.331 | 0.145 | 0.218 | 0.958 | 0.883 |
| 7 | 0 | 0.916 | 0.797 | 0.205 | 0.326 | 0.145 | 0.210 | 0.958 | 0.893 |
| | 1 | 0.919 | 0.805 | 0.202 | 0.318 | 0.143 | 0.209 | 0.959 | 0.898 |
| | 2 | 0.915 | 0.803 | 0.205 | 0.324 | 0.146 | 0.210 | 0.957 | 0.896 |
| | 3 | 0.918 | 0.793 | 0.205 | 0.306 | 0.145 | 0.200 | 0.959 | 0.891 |
| | 4 | 0.918 | 0.773 | 0.204 | 0.335 | 0.143 | 0.219 | 0.959 | 0.879 |

**c**

**Average prediction model performance**

| Feature set | Model performace metrics (Average of 5-fold cross-validation results) | | | | | | | |
|---|---|---|---|---|---|---|---|---|
| | $R^2$ | | RMSE | | MAE | | Pearson r | |
| | Train | Test | Train | Test | Train | Test | Train | Test |
| 5 | 0.884 | 0.773 | 0.242 | 0.338 | 0.166 | 0.218 | 0.941 | 0.880 |
| 6A | 0.896 | 0.793 | 0.229 | 0.323 | 0.158 | 0.210 | 0.947 | 0.891 |
| 6B | 0.915 | 0.789 | 0.206 | 0.326 | 0.146 | 0.212 | 0.958 | 0.889 |
| 7 | 0.917 | 0.794 | 0.204 | 0.322 | 0.144 | 0.209 | 0.958 | 0.891 |

**d**

**Statistical analysis**

| Comparison | Paired t-test (*p*-values) | | | | | | | |
|---|---|---|---|---|---|---|---|---|
| | $R^2$ | | RMSE | | MAE | | Pearson r | |
| | Train | Test | Train | Test | Train | Test | Train | Test |
| 5 vs 6A | 0.0881 | 0.0033 | 0.0826 | 0.0034 | 0.1009 | < 0.0001 | 0.0841 | 0.0025 |
| 5 vs 6B | < 0.0001 | 0.0081 | < 0.0001 | 0.0064 | < 0.0001 | 0.0079 | < 0.0001 | 0.0077 |
| 5 vs 7 | < 0.0001 | 0.0204 | < 0.0001 | 0.0190 | < 0.0001 | 0.0054 | < 0.0001 | 0.0174 |

**Extended Data Fig. 5 Machine learning-based model training identifies optimal structural feature set for Detectron performance prediction. a**, Schematic of the machine learning workflow using XGBoost regression trained on the indicated feature sets. **b**, Model performance for each fold was evaluated using coefficient of determination ($R^2$), root mean squared error (RMSE), mean absolute error (MAE) and Pearson's correlation coefficient (*r*). **c**, Average of performance metrics across the five outer cross-validation folds. **d**, Paired *t*-tests comparing performance metrics between the indicated feature-set models.

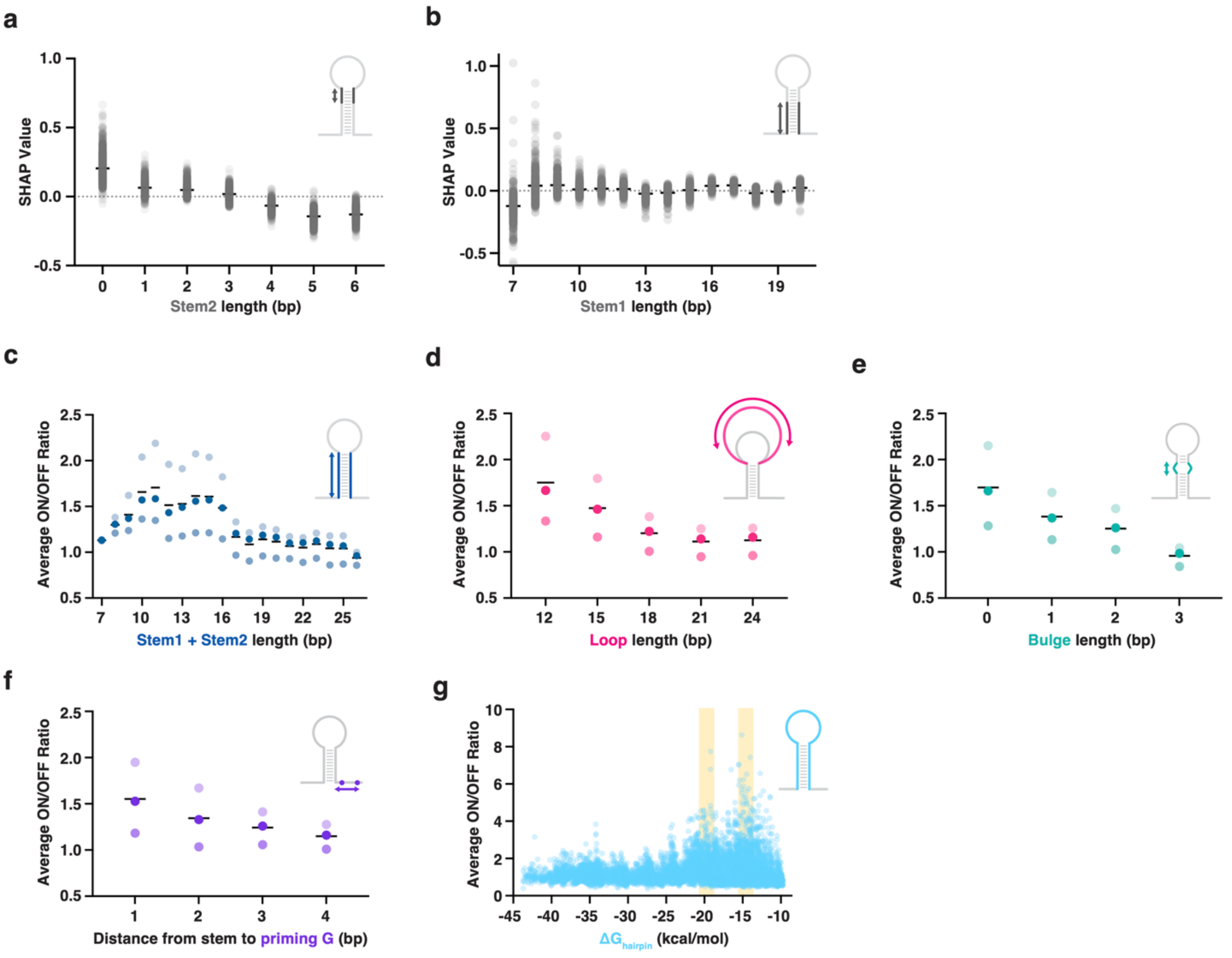


**Extended Data Fig. 6 Average ON/OFF ratios and SHAP feature effects across Detectron structural parameters. a-b**, SHAP analysis of ON/OFF ratios across (a) stem2 and (b) stem1 lengths. Each circle represents an individual Detectron variant. Bars represent the mean SHAP value of the variants. **c-f**, Average ON/OFF ratios across different feature distributions: (c) combined stem lengths; (d) loop lengths; (e) bulge lengths; and (f) distance from stem to priming G. Closed circles indicate three biological replicates. Bars represent the mean (±s.d.) of three biological replicates. **g**, Average ON/OFF ratios plotted against free energy of the hairpin ($\Delta G_{hairpin}$). Each circle represents an individual Detectron variant.

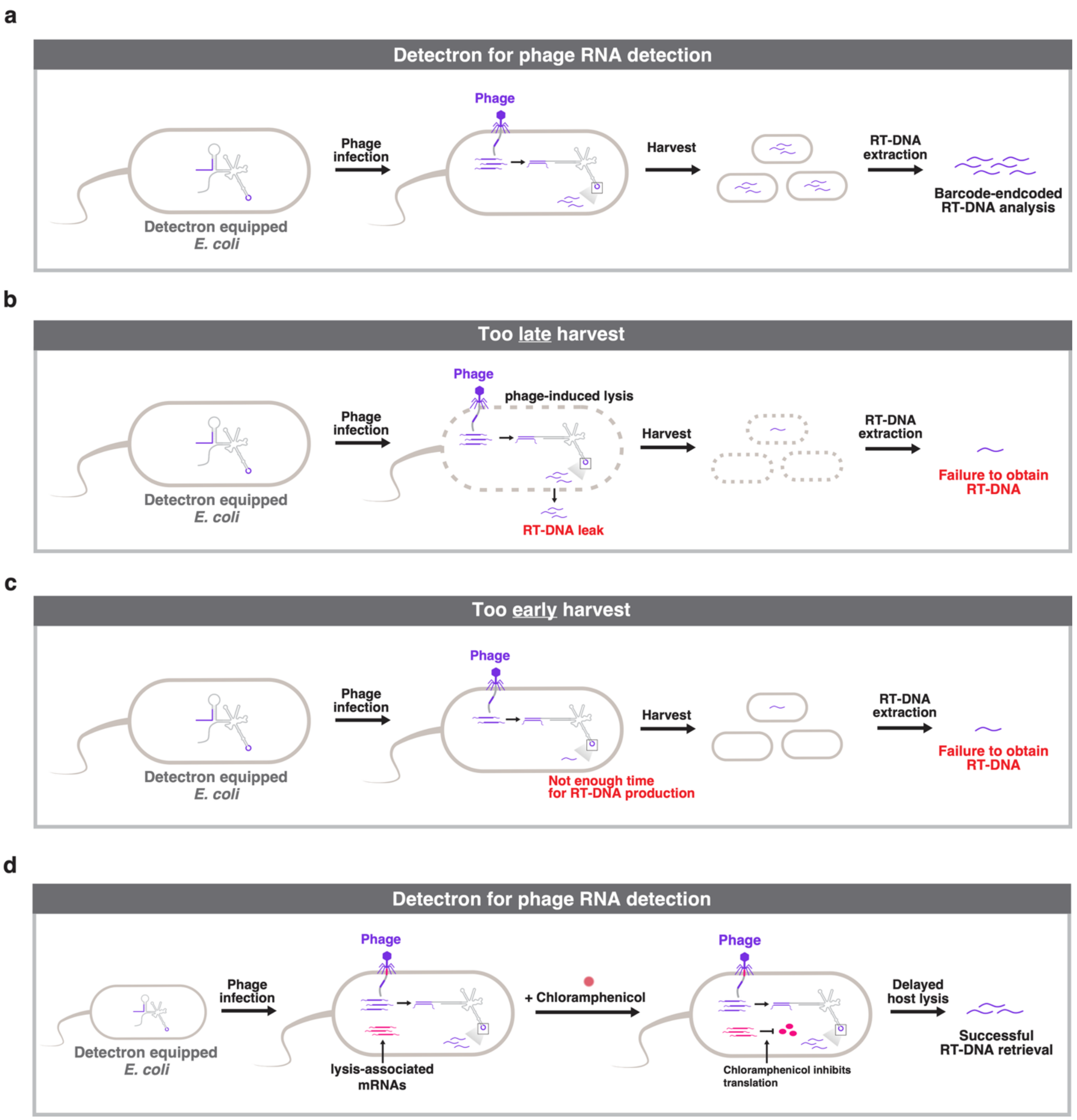


**Extended Data Fig. 7 Chloramphenicol treatment for Detectron-mediated detection of phage infection.** **a**, Schematic overview of Detectron-mediated detection of phage-derived mRNAs during infection. **b**, Late harvest scenario. Prolonged incubation after infection leads to phage-induced host lysis and leakage of RT-DNA into the medium, resulting in loss of detectable barcodes. **c**, Early harvest scenario. Premature sampling fails to allow sufficient time for RT-DNA production, yielding weak or undetectable barcode signal. **d**, Optimized condition with chloramphenicol treatment. Addition of chloramphenicol post-infection inhibits translation of lysis-associated mRNAs, thereby delaying host lysis while allowing continued accumulation of intracellular barcode-encoded RT-DNA.
